# *Rhodosporidium toruloides:* A new platform organism for conversion of lignocellulose into terpene biofuels and bioproducts

**DOI:** 10.1101/154872

**Authors:** Junko Yaegashi, James Kirby, Masakazu Ito, Jian Sun, Tanmoy Dutta, Mona Mirsiaghi, Eric R. Sundstrom, Alberto Rodriguez, Edward Baidoo, Deepti Tanjore, Todd Pray, Kenneth Sale, Seema Singh, Jay D. Keasling, Blake A. Simmons, Steven W. Singer, Jon K. Magnuson, Adam P. Arkin, Jeffrey M. Skerker, John M. Gladden

## Abstract

**Background:** Economical conversion of lignocellulosic biomass into biofuels and bioproducts is central to the establishment of a robust bioeconomy. This requires a conversion host that is able to both efficiently assimilate the major lignocellulose-derived carbon sources and divert their metabolites toward specific bioproducts.

**Results:** In this study, the carotenogenic yeast *Rhodosporidium toruloides* was examined for its ability to convert lignocellulose into two non-native sesquiterpenes with biofuel (bisabolene) and pharmaceutical (amorphadiene) applications. We found that *R. toruloides* can efficiently convert a mixture of glucose and xylose from hydrolyzed lignocellulose into these bioproducts, and unlike many conventional production hosts, its growth and productivity were enhanced in lignocellulosic hydrolysates relative to purified substrates. This organism was demonstrated to have superior growth in corn stover hydrolysates prepared by two different pretreatment methods, one using a novel biocompatible ionic liquid (IL) choline α-ketoglutarate, which produced 261 mg/L of bisabolene at bench-scale, and the other using an alkaline pretreatment, which produced 680 mg/L of bisabolene in a high gravity fed-batch bioreactor. Interestingly, *R. toruloides* was also observed to assimilate *p-*coumaric acid liberated from acylated grass lignin in the IL hydrolysate, a finding we verified with purified substrates. *R. toruloides* was also able to consume several additional compounds with aromatic motifs similar to lignin monomers, suggesting that this organism may have the metabolic potential to convert depolymerized lignin streams alongside lignocellulosic sugars.

**Conclusions:** This study highlights the natural compatibility of *R. toruloides* with bioprocess conditions relevant to lignocellulosic biorefineries and demonstrates its ability to produce non-native terpenes.

## Background

Growing energy demands and concerns over global warming and environmental pollution associated with the consumption of petroleum have made it imperative to develop and foster a bioeconomy focused on efficient low-carbon emission technologies. Replacement of petroleum-derived fuels and chemicals with bio-based alternatives derived from renewable carbon sources has been identified as a promising approach to help realize a bioeconomy, and provides an added benefit of supporting domestic economies through localized revenues and job growth [1]. Lignocellulosic biomass, composed primarily of cellulose, hemicellulose and lignin, is the most abundant renewable carbon source available today, and has been widely studied as a substrate for microbial production of bio-based fuels and chemicals. Most of these efforts have focused on converting one or two of the major components of plant biomass, primarily cellulose and hemicellulose, but none of them have demonstrated conversion of all three components by a single microbe into a single non-native bioproduct. Due to its heterogeneity and recalcitrance to depolymerization, the cross-linked phenolic polymer lignin is the most underutilized of the three components for bioconversion, and is often relegated to being burned for heat and energy generation in a biorefinery [2]. However, technoeconomic (TEA) and life-cycle (LCA) analyses have indicated that lignin valorization will be critical for maintaining the economic viability and sustainability of lignocellulosic biorefineries [3]. Therefore, conversion strategies that incorporate lignin will have an economic advantage over those that focus solely on carbohydrate conversion, creating a strong incentive for biorefineries to adopt microbial conversion hosts that can achieve this feat.

While well-established microbes such as *Escherichia coli* and *Saccharomyces cerevisiae* are convenient hosts for bioproduct synthesis from glucose or xylose, they do not readily utilize multiple carbon sources simultaneously, especially not those derived from lignin, making it difficult to efficiently use lignocellulose as a carbon source [4]. Two approaches to circumvent this problem are to 1) engineer commonly used hosts such as *E. coli* and *S. cerevisiae* to efficiently utilize cellulose, hemicellulose, and lignin depolymerization products, or 2) find a host that naturally has this ability and engineer it to make bioproducts. *Rhodosporidium toruloides*, an oleaginous, carotenogenic basidiomycete yeast, has been studied as a model organism for lipid production and has been shown to co-utilize both hexose and pentose sugars [5], suggesting potential advantages of *R. toruloides* over conventional lignocellulosic conversion hosts. *R. toruloides* accumulates high concentrations of lipids and carotenoids, both of which are derived from acetyl-CoA [6]. This suggests that it may be a promising host for the production of compounds synthesized from acetyl-CoA, especially terpene and lipid-based bioproducts. Not only does it make these natural bioproducts, it can also grow to very high cell densities (100 g/L dry cell mass) [7], another important industrially-relevant characteristic. Taking advantage of the recently developed genetic tools for *R. toruloides* [8-11], we explored its utility as a new platform for production of non-native terpenes from lignocellulose. We demonstrate that *R. toruloides* has the unique ability to simultaneously utilize glucose and xylose derived from depolymerized cellulose and hemicellulose in addition to compounds associated with lignin, such as *p*-coumaric acid. This compound is associated with grass lignins (like corn, sorghum, switchgrass, etc.) where it is conjugated to lignin through an ester linkage that is easily cleavable using existing alkaline lignocellulosic pretreatment technologies [12]. Unlike this lignin conjugate, the lignin polymer itself is very heterogeneous and its depolymerization has the potential to yield a wide variety of compounds derived from the lignin *p*-hydroxyphenyl (H), guaiacyl (G), and syringyl (S) phenylpropanoid units. Therefore, *R. toruloides* was also examined for its ability to consume several compounds containing aromatic motifs similar to the H, G and S lignin subunits. Demonstration of the conversion of these types of compounds along with lignocellulosic sugars opens the possibility of including a lignin as a carbon source in lignocellulose conversion schemes, a process that would increase the efficiency and commercial viability of a biorefinery. Finally, we demonstrate that *R. toruloides* is compatible with a single-unit or one-pot lignocellulose pretreatment, saccharification, and fermentation process (Fig. 1) that potentially reduce biorefinery capital and operating expenses (CAPEX, and OPEX, respectively) and wastewater treatment [13, 14]. Together, these abilities suggest that *R. toruloides* has the potential to be a platform organism for the conversion of the majority of the carbon present in lignocellulose into advanced biofuels and bioproducts.

**Fig. 1.**
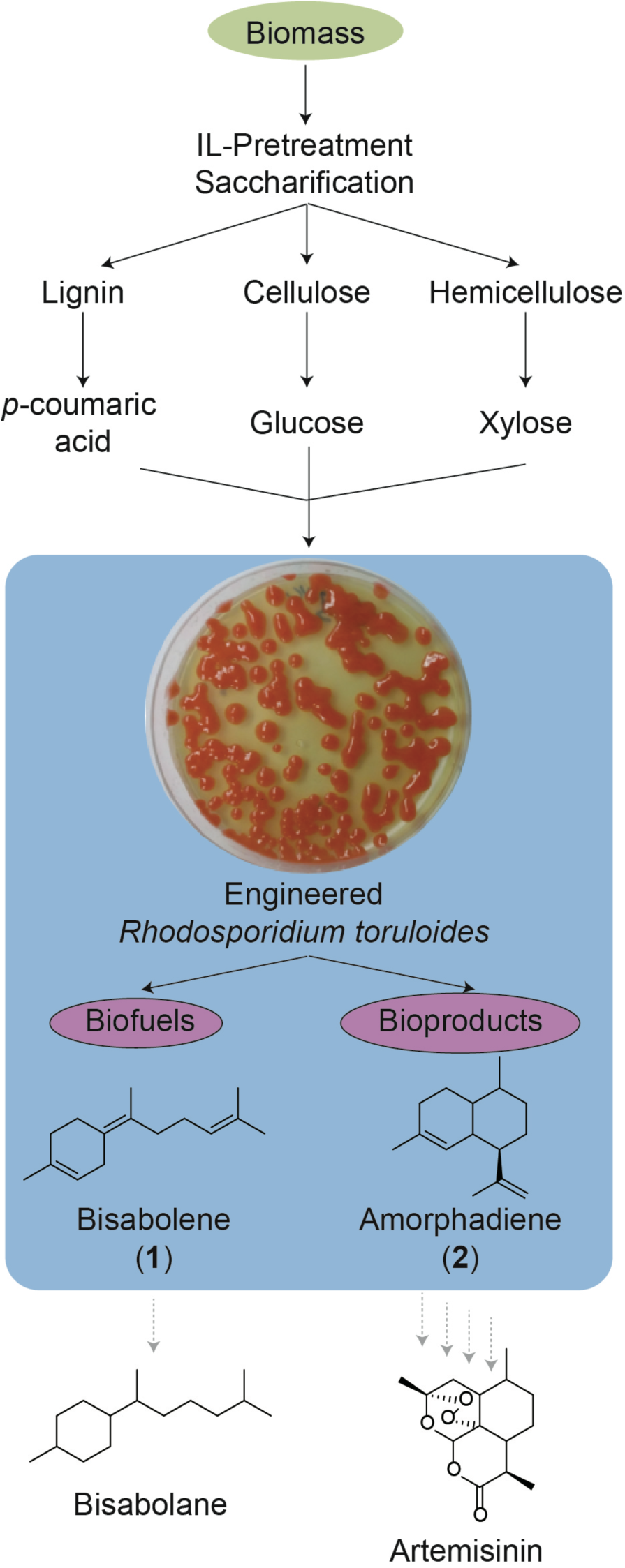
Schematic of *R. toruloides* as a new platform for production of lignocellulosic biofuels and bioproducts.

## Methods

### Media

Synthetic defined (SD) medium was made following manufacturers’ instructions with Difco yeast nitrogen base (YNB) without amino acids (Becton, Dickinson & Co., Sparks, MD) and Complete Supplemental Mixture (CSM; Sunrise Science Products, San Diego, CA). Initial medium pH was adjusted to 7.4 with NaOH unless otherwise stated. Luria Broth (LB) and Yeast Peptone Dextrose (YPD) media were made using pre-mixed Difco LB broth and Difco YPD broth, respectively.

### Growth conditions

*R. toruloides* seed cultures were obtained by inoculating 5 mL LB with single colonies from a YPD agar plate containing antibiotics at the following concentrations: nourseothricin, 100 μg/mL, and cefotaxime, 300 μg/mL. The seed cultures were used to inoculate 5 mL SD media with a starting optical density at 600 nm (OD_600_) of 0.1. The same inoculation strategy was used for cultivations of lignin-related monoaromatics, with the following compounds being added separately to SD medium: *p*-coumaric acid, ferulic acid, *p*-hydroxybenzoic acid, vanillic acid, sinapic acid, benzoic acid and vanillin, at a final concentration of 2 g/L for each compound. Cultures of terpene-producing strains were overlaid with 20% (v/v) dodecane. All cultures were grown at 30°C with shaking at 200 rpm. Growth was monitored by measuring OD_600_. Samples in which the OD measurements were significantly different from others in the sample set were excluded from the analysis.

### Plasmid construction and transformation

Strains and plasmids used in this study can be found in Table 1, and are also available through the Joint BioEnergy Institute Strain Registry <u>(https://public-registry.jbei.org/</u> [15]) and are available upon request.

**Table 1.**
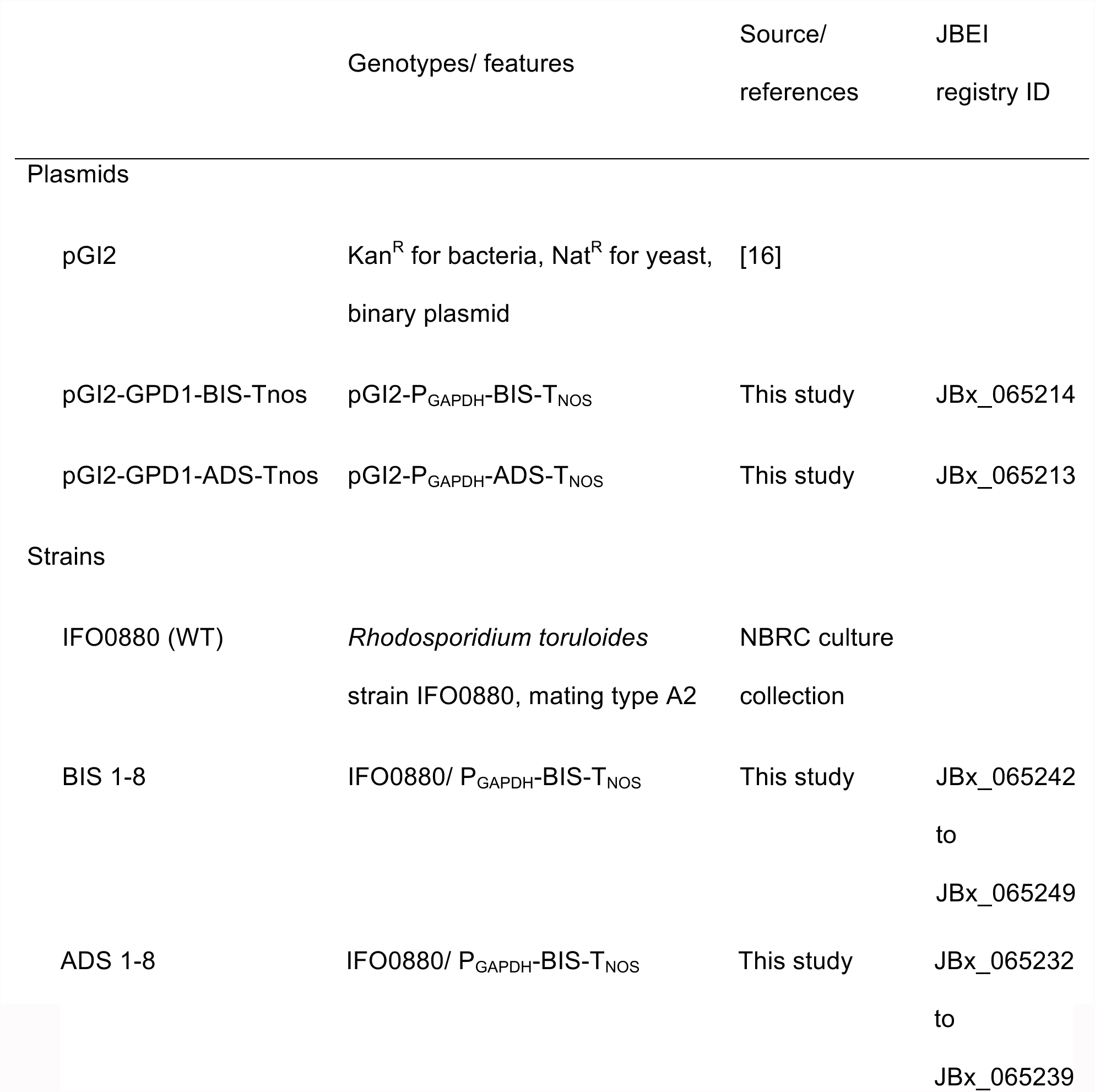
Strains and plasmids used in this study.

Codon optimization, gene synthesis, and plasmid construction were performed by GenScript (Piscataway, NJ). The genes encoding bisabolene synthase (*BIS*) and amorphadiene synthase (*ADS*) were codon optimized for *R. toruloides* based on a custom IFO0880 codon usage table http://genome.jgi.doe.gov/Rhoto_IFO0880_3/Rhoto_IFO0880_3.home.html), and the constructs were designed so that each gene was positioned between the *GAPDH* promoter and *NOS* terminator [10]. The constructs were synthesized and inserted into the *Agrobacterium tumefaciens*-mediated transformation (ATMT) plasmid pGI2 [16] using the EcoRV restriction sites. The pGI2-derived plasmids were introduced into *R. toruloides* recipient strains by ATMT as previously described [8, 10].

### Measurement of bisabolene and amorphadiene

For measurement of bisabolene production, 10 μL of the dodecane overlay was sampled and diluted into 390 μL of ethyl acetate spiked with 1 mg/L caryophyllene as an internal standard. Bisabolene was quantified by gas chromatography-mass spectrometry (GC-MS) as described previously [17].

### Assessment of genetic stability of the bisabolene-producing strain BIS3

Strain BIS3 was cultured overnight in LB and, after removal of medium, cells were used to inoculate 5 mL of SD medium containing 2% (w/v) glucose at a starting OD_600_ of 0.1, in triplicate. Cultures were overlaid with 20% dodecane and incubated by shaking at 200 rpm at 30°C. After 6 days the cultures were used to inoculate fresh medium of the same type and dodecane was sampled to quantify bisabolene production. This process was repeated 2 additional times, spanning four rounds of culture over 24 days.

### Measurement of lipid and carotenoid content

Total lipid content was quantified gravimetrically following extraction with Folch reagent (2:1 chloroform/methanol) as described previously [18]. Carotenoids were extracted with acetone and quantified by high performance liquid chromatography (HPLC) as described previously [19]. Carotenoid standards β-carotene, torulene, and torularhodin were obtained from Carotenature, GmbH (Ostermundigen, Switzerland).

### Analysis of sugars and α-ketoglutarate

The concentrations of sugars and α-ketoglutarate were quantified on an Agilent Technologies 1200 series HPLC equipped with an Aminex HPX-87H column (BioRad, Hercules, CA) as described previously [20]. Sugars were monitored by a refractive index detector, and α-ketoglutarate was monitored by a diode array detector at 210 nm. Concentrations were calculated by integration of peak areas and comparison to standard curves for the compounds of interest.

### Analysis of monomeric aromatics

The concentrations of monoaromatics were quantified on an Agilent Technologies 1200 series HPLC equipped with a Zorbax Eclipse XDB-C18 column (Agilent Technologies, Santa Clara, CA). Five microliters of each sample was injected onto the column and eluted isocratically with 20% (v/v) acetonitrile in H_2_O containing 0.5% acetic acid (v/v) at a flow rate of 1 mL/min for 15 minutes at a column temperature of 24°C. Detection of *p*-coumaric acid and sinapic acid utilized a diode array detector at 310 nm, while ferulic acid, *p*-hydroxybenzoic acid, vanillic acid, benzoic acid and vanillin utilized a diode array detector at 275 nm. Concentrations were calculated by comparison of peak areas to calibration curves generated with high purity standards.

### Preparation of the ionic liquid [Ch][α-Kg]

In a typical process, [Ch][OH] (46 wt% in H_2_O) was mixed with α-ketoglutaric acid (40 wt% in H_2_O) in a 2:1 weight ratio at room temperature. The pH of the resulting ionic liquid (IL) was measured by a pH probe and maintained at 13.5.

### Ionic liquid one-pot pretreatment and saccharification

Cellulase (Cellic^®^ CTec2; Batch# VCN10001, protein content 188 mg/mL) and hemicellulase (Cellic^®^ HTec2; Batch# VHN00001, protein content 180 mg/mL) enzyme mixtures were received as gifts from Novozymes NA (Franklinton, NC, USA), and mixed with a volume ratio of 9:1 before use. Corn stover was supplied by the Department of Chemical Engineering & Materials Science at Michigan State University. The biomass was ground by a Thomas-Wiley Mini Mill fitted with a 20-mesh screen (Model 3383-L10 Arthur H. Thomas Co., Philadelphia, PA, USA) and analyzed for polysaccharide composition after drying (glucan 38.90%±0.04, xylan 24.77 wt%±0.01, and lignin 18.42 wt%±0.27).

In an integrated process, 40 g of corn stover was mixed with 160 g [Ch][α-Kg] (40 wt% in H_2_O) at a 20 wt% biomass loading in a 300 mL Parr reactor and pretreated at 120°C for 4 h. After pretreatment, the slurry was diluted with deionized (DI) water to obtain a final IL concentration of 10 wt%. Before adding the enzyme mixture (CTec2/HTec2=9:1, v/v) for the saccharification, α-ketoglutaric acid (40 wt% in H_2_O) was used to lower the pH of the system to 5. Enzymatic hydrolysis was conducted at 50°C in a 1 L shake flask for 3 days with an enzyme loading of 20 mg protein/g corn stover. Titers were: glucose (17.1 g/L), xylose (9.1 g/L), and *p*-coumaric acid (383 mg/L).

The raw and pretreated corn stover were dried and characterized with powder X-ray diffraction (XRD). The XRD analyses were performed on a PANalytical Empyrean X-ray diffractometer equipped with a PIXcel3D detector and operated at 45 kV and 40 kA using Cu Kα radiation (λ=1.5418Å). The patterns were collected in the 2θ range from 5 to 60° with a step size of 0.039° and an exposure time of 300 seconds. A reflection-transmission spinner was used as a sample holder and the spinning rate was set at 8 rpm throughout the experiment. Crystallinity index (CrI) was determined by Segal’s method [21].

X-ray diffraction (XRD) studies were conducted to determine the changes in the crystalline vs. non-crystalline components found in the untreated corn stover, and to monitor the structural changes in these polymers that occur during the pretreatment process. Additional file 1 shows the X-ray diffractograms of the untreated and pretreated corn stover after processing at 120°C for 4 h. The diffractogram obtained from the untreated corn stover has two major diffraction peaks at 22.5° and 15.7° 2θ, characteristic of the cellulose I polymorph that corresponds to [002] and combined [101] + [10^−^1] lattice planes, respectively. The third small peak at 34.5° ([040] lattice plane) corresponds to 1/4 of the length of one cellobiose unit and arises from ordering along the fiber direction [21, 22]. The diffractogram obtained from pretreated corn stover still retains the cellulose I polymorph, with a relative decrease in the intensity of the [002] peak. The crystallinity index (CrI) of corn stover decreased from 61% to 50% after pretreatment. This decrease in the cellulose crystallinity after pretreatment is reflected in the high saccharification efficiency observed.

### Alkaline biomass pretreatment and saccharification for bioreactor cultivation

A mixture containing 15% corn stover biomass (7 wt% moisture), 1.5% NaOH, and 83.5% water was pretreated by autoclave at 121°C for 1 h. Following pretreatment, the biomass was wrapped in cheesecloth and dried in a laundry centrifuge to approximately 30 wt% solids. The supernatant was discarded and biomass was re-suspended and soaked in DI water overnight after the pH was adjusted to 5.0. The pH-adjusted biomass was then centrifuged a second time to remove excess salt and moisture.

Pretreated biomass, containing 85% w/w moisture, was saccharified in 2 L IKA reactors (model LR-2.ST, IKA, USA) using commercially available enzymes CTec2 and HTec2 (Novozymes, USA). Enzymes with the following loadings were added to the reactor: 64 mg CTec2/g dry biomass and 8 mg HTec2/g dry biomass. Enzymatic saccharification was performed at 50°C with pH in the range of 4.5 to 5.5 for 96 h. Upon completion of the saccharification reaction, the unhydrolyzed biomass was separated from the hydrolysate by centrifugation at 4000 x g for 30 min. The hydrolysate was filtered with 0.7 μm and then 0.45 μm filter papers to separate any remaining particles and finally sterilized by passing through 0.2 μm filters and stored at 4°C until use. The final hydrolysate contained 86.5 g/L of free glucose and 38.1 g/L of free xylose.

### Bioreactor cultivation using alkaline hydrolysate

The seed cultures were prepared by transferring a single colony from a YPD agar plate to a 500 mL baffled flask containing 250 mL of seed medium. The seed medium consisted of 10 g/L yeast extract, 20 g/L peptone, and 20 g/L glucose. The seed was grown at 30°C, shaking at 250 rpm overnight to reach exponential growth phase. When the seed reached the exponential growth phase, 5.5% (v/v) inoculum was transferred to each bioreactor to reach an initial OD_600_ of 0.6. Bisabolene production in *R. toruloides* was examined in 2 L bioreactors (BIOSTAT B, Sartorius, Germany) with an extractive fermentation. Fermentation process parameters were controlled with temperature at 30°C, dissolved oxygen at 40% air saturation, and pH 5, respectively. Dissolved oxygen was controlled by adjusting the agitation rate at a constant airflow. Culture pH was controlled at 5 by automated addition of 2 M NaOH. Foaming was controlled by addition of 5% (v/v) Antifoam 204 as needed.

The batch medium in the fermenter containing SD medium had the following concentrations: 20 g/L glucose, 6.7 g/L YNB, and 0.79 g/L CSM. The feed for this reactor during fed-batch growth was 500 g/L glucose in DI water. The batch medium for the reactor containing the alkaline hydrolysate had the following concentration: 20% (v/v) corn stover alkaline hydrolysate, 6.7 g/L YNB, and 0.79 g/L CSM. The concentration of glucose and xylose in the batch medium for this reactor was 17.3 and 7.62 g/L, respectively. The feed for the fed-batch phase for this reactor was corn stover alkaline hydrolysate containing 86.5 g/L of glucose and 38.1 g/L of xylose. Both fermenters contained 700 mL of initial batch medium and 150 mL of dodecane overlay containing 1 g/L of pentadecane as internal standard to account for evaporation of the overlay. The feed was initiated once all the sugar in the batch medium was consumed. The feed flow rate was adjusted to maintain glucose concentration in the reactors below 5 g/L. Cell growth and bisabolene production was monitored by taking 5 mL samples at predetermined time points.

### Dry cell weight analysis of bioreactor cultivations

For dry cell weight analysis 10 mL sample was removed from each reactor. After separation of the aqueous and organic phase, 5 mL sample from the aqueous phase was transferred to pre-weighed falcon tubes. The cells were centrifuged at 4000 x g for 10 min and supernatant was discarded. Cells were dried in a vacuum oven (Binder, Germany) at 60°C until the weight was stable.

## Results and discussion

### *R. toruloides* as a platform for terpene production

Terpenes are produced by a variety of organisms and have a wide range of applications from flavors, fragrances, and pharmaceuticals to biofuels and chemical feedstocks [23]. In this study, we selected two terpenes, amorphadiene and bisabolene, to examine the suitability of *R. toruloides* as a lignocellulosic conversion host. Amorphadiene, a precursor of the antimalarial drug artemisinin, was chosen as an example of a commercially relevant bioproduct [24] and bisabolene, an immediate precursor of the D2 diesel alternative bisabolane, was chosen as an example of an advanced “drop-in” biofuel [25]. Codon optimized expression cassettes for bisabolene (BIS) and amorphadiene (ADS) synthases were constructed and separately integrated into the genome of *R. toruloides* IFO0880 using *Agrobacterium tumefaciens* mediated transformation (ATMT) [8]. A number of transformants were confirmed to produce either bisabolene (Fig. 2A) or amorphadiene (Fig. 2B), with variance in titer between strains most likely due to copy number and integration site effects [26, 27]. Terpene titers for selected strains in synthetic defined (SD) medium containing 2% (w/v) glucose, reached 294 mg/L for bisabolene and 36 mg/L for amorphadiene. These bisabolene and amorphadiene titers attained in *R. toruloides* are highly encouraging considering that they exploit the natural flux of carbon through this yeast’s native terpene biosynthetic pathway. In comparison, the yeast *S. cerevisiae* transformed with high copy plasmids harboring the BIS and ADS genes and grown in equivalent media attained significantly lower bisabolene and amorphadiene titers – approximately 20 and 10 mg/L, respectively ([28] for ADS, unpublished data for BIS). Another notable feature of the *R. toruloides* BIS strain is that bisabolene titers show remarkable stability over extended periods of serial cultivation, varying by less than 16% over the course of four cultures spanning 24 days (Fig. 3). It should be noted that this reproducibility was also achieved without the need for a heterologous inducer or antibiotic selection, since the BIS gene is stably integrated into the genome and its expression is under control of a constitutive GAPDH promoter [8-10]. Both of these features reduce OPEX in a biorefinery. In comparison, the bisabolene titer from an engineered strain of *S. cerevisiae* grown under similar conditions was found to decline by more than 75% over 14 days [29]. The strain stability we observed in engineered *R. toruloides* is an important industrial phenotype and a critical factor for large-scale economical production of any bioproduct.

**Fig. 2.**
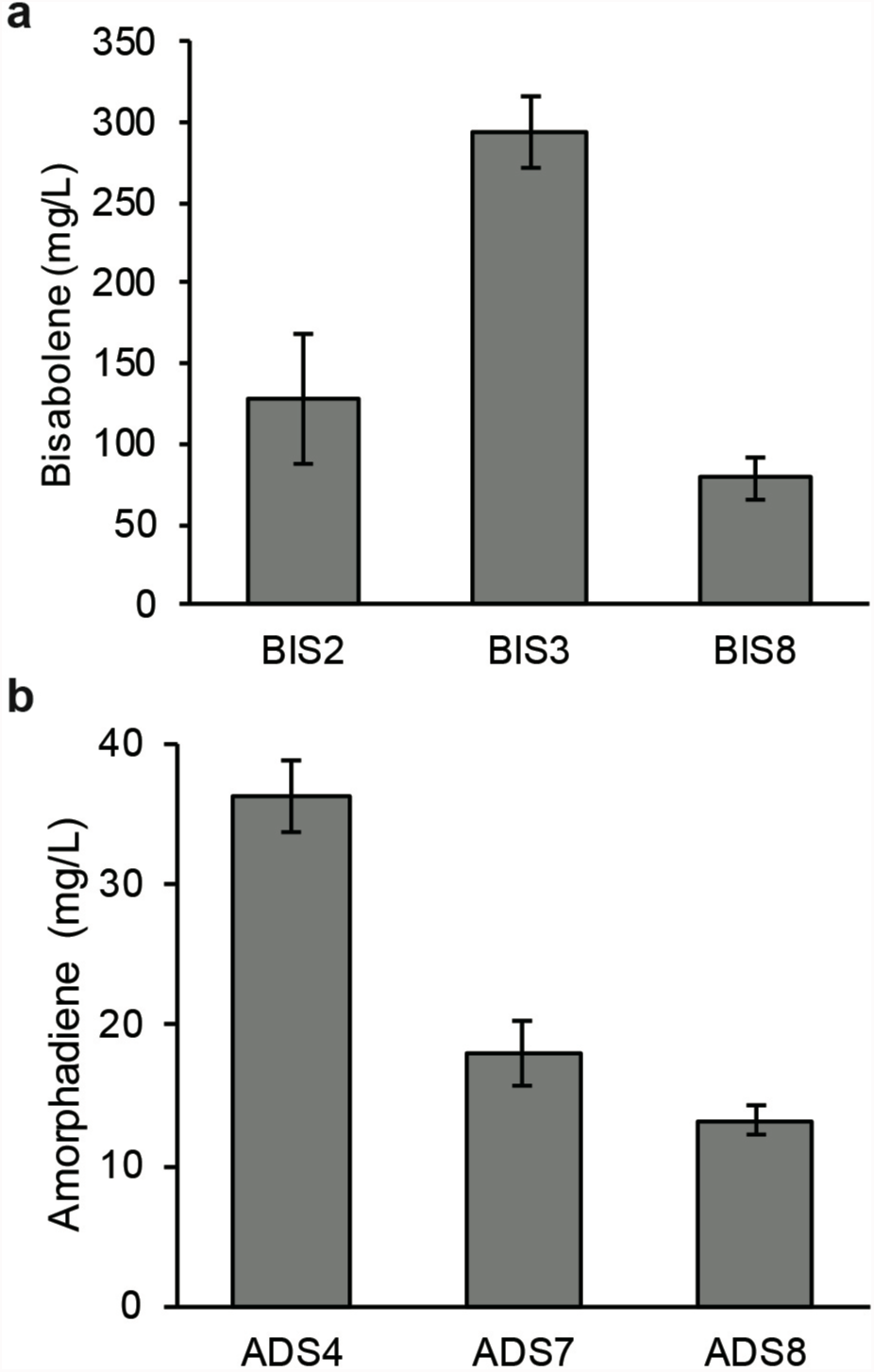
Terpene titers of *R. toruloides* transformants. (**A**) Bisabolene and (**B**) amorphadiene titers in selected strains grown in SD medium with 2% glucose. 5 mL cultures in test tubes were set up at a starting OD of 0.1 with a 20% dodecane overlay. At day 7, the dodecane layer was sampled and analyzed for bisabolene or amorphadiene. (n=3, data shown as average ±s.d, representative from two independent experiments).

**Fig. 3.**
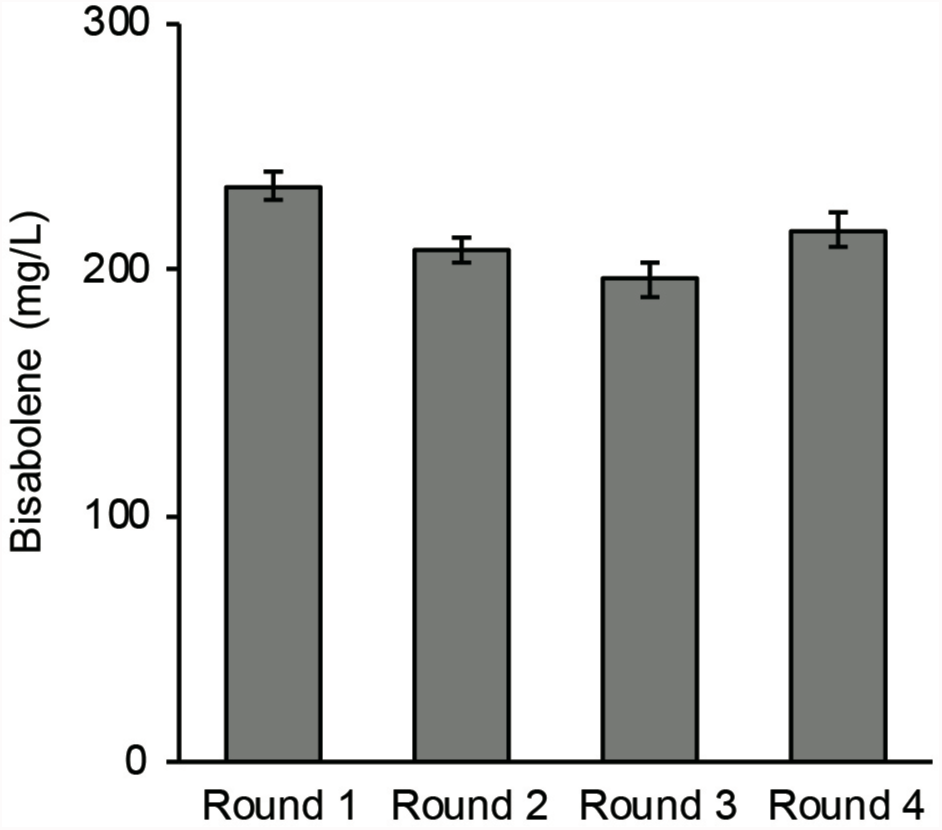
Stability of bisabolene production in serial cultures. Cultures in SD medium with 2% glucose were passaged consecutively every 6 days. (n=3, data shown as average ±s.d, from a single experiment).

We found that the pH of the growth medium is an important factor for efficient sugar utilization by *R. toruloides*. After examining a range of starting pH values in unbuffered medium (3 to 8) in batch cultures, a starting pH of 7.4 was determined to be optimal to achieve complete glucose utilization (Table 2) and the highest bisabolene titer (Fig. 4). Interestingly, *R. toruloides* grew and produced bisabolene at a pH as low as 3.4, suggesting the host may be amenable to production of organic acids or other bioproducts that require low pH. One potential explanation for the decline in pH is that *R. toruloides* is producing native organic acids of potential value, a topic that merits further investigation. However, once the pH declines to 2.5 (in unbuffered medium starting at pH 7 or below), sugar utilization is strongly inhibited, suggesting that the pH must remain above this level to enable efficient carbon conversion. Therefore, all subsequent experiments in unbuffered media were performed with a starting pH of 7.4.

**Table 2.** Percent utilization of glucose in SD media starting at various pH.

| pH | % Utilization |
| --- | --- |
| 3.4 | 52.6 ± 5.4 |
| 4.4 | 67.0 ± 4.7 |
| 5.4 | 69.3 ± 3.5 |
| 6.4 | 76.4 ± 1.3 |
| 7.4 | 100.0 ± 0.0 |
| 8.4 | 96.8 ± 2.4 |

Cultures were carried out as described above, the aqueous layer was sampled for glucose analysis (n=3, data shown as average ±s.d, from a single experiment).

**Fig. 4.**
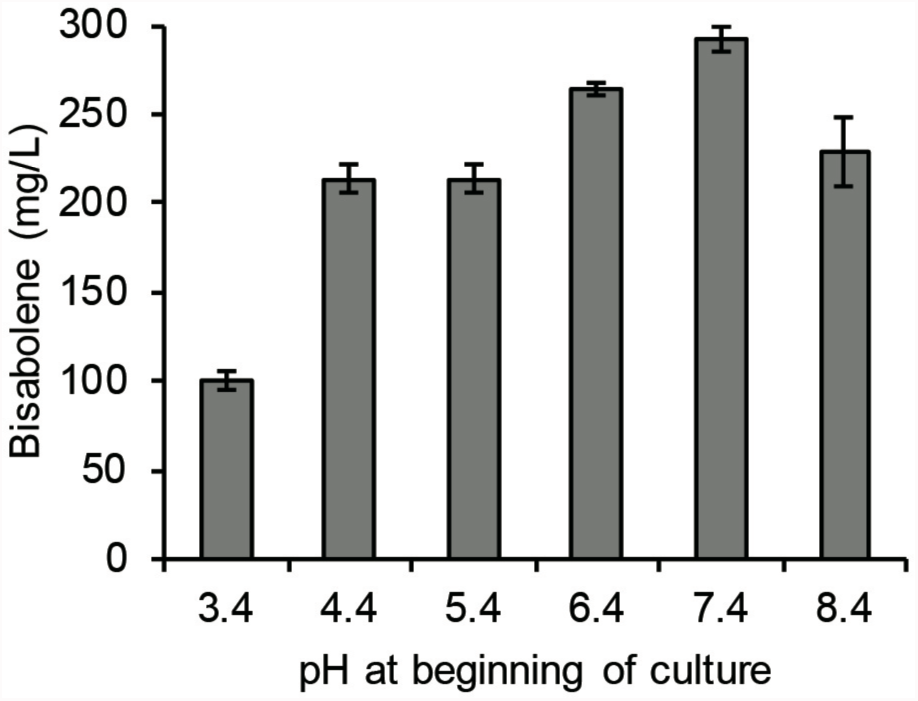
The effect of pH on bisabolene titers. (n=3, data shown as average ±s.d, from a single experiment in SD medium with 2% glucose).

### *R. toruloides* can convert multiple carbon sources into bisabolene in defined media

To demonstrate the capability of engineered *R. toruloides* to utilize different carbon sources to produce non-native terpenes, we cultivated the bisabolene-producing strain BIS3 with the most abundant sugars present in lignocellulosic hydrolysates: glucose and xylose. In addition, we observed the liberation and consumption of *p*-coumaric acid in the IL-pretreated cornstover hydrolysate described in detail below (the hydrolysate in Fig. 7), and therefore we examined this compound as a carbon source along with the sugars. This phenolic compound is associated with grass lignins through an ester linkage to lignin monomers formed prior to lignin polymerization [12]. This ester linkage can easily be cleaved under alkaline conditions [30], like those found in the pretreatments used to generate hydrolysates in this study. Initially, these three carbon sources were provided individually and growth, carbon utilization, and bisabolene production were monitored (Fig. 5A-C). Glucose was completely consumed at the fastest rate, followed by *p-*coumaric acid, then xylose (in 1, 3, and 4 days, respectively). The highest bisabolene titers were observed in the *p*-coumaric acid cultures, likely due to its higher percentage of carbon relative to the sugars (Fig. 5A-C). Remarkably, when combined, all three carbon sources were completely utilized within four days (Fig. 5D). The *p-*coumaric acid was completely utilized earlier in the presence of the other sugars (2 vs 3 days), while complete utilization of glucose and xylose took slightly longer when present in the mixture (glucose: 2 vs 1 day; xylose: both day 4 but less consumed by day 2 in the mixture).

**Fig. 5.**
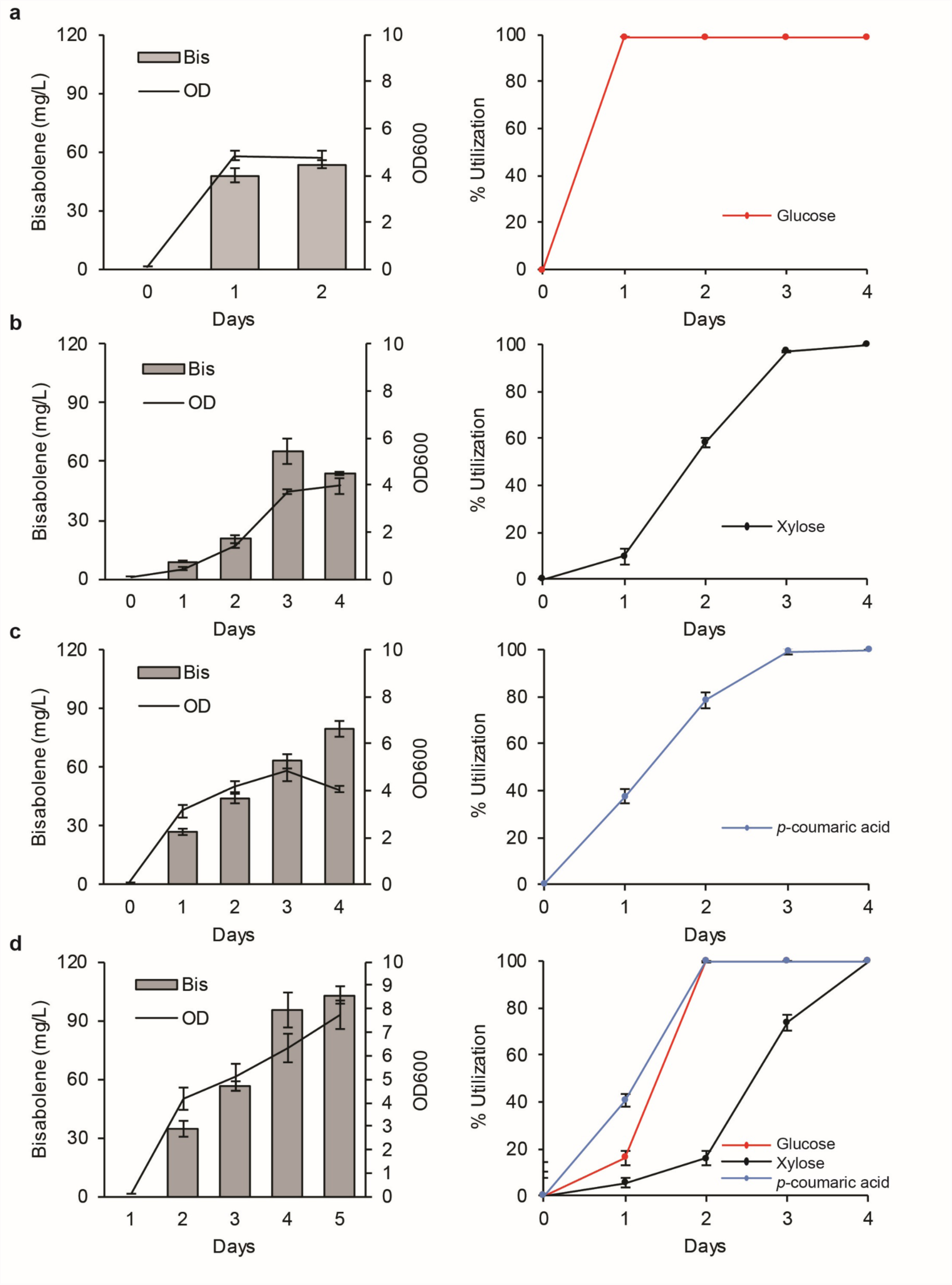
Conversion of glucose, xylose, and *p-*coumaric acid, both individually and mixed, into bisabolene by *R. toruloides*. Bisabolene titers, growth, and carbon utilization of strain BIS3 grown in SD medium supplemented with different carbon sources: (**a**) 0.5% Glucose, (**b**) 0.5% xylose, (**c**) 0.5% *p-* coumaric acid, and (**d**) 0.5% glucose, 0.5% xylose, 0.5%*p-*coumaric acid. Left panels: Lines represent ODs, bars represent bisabolene titers. Right panels: glucose (red), xylose (black), *p-*coumaric acid (blue). 5 mL cultures in test tubes were set up at a starting OD of 0.1 with a 20% dodecane overlay. At each time point, the dodecane layer was sampled and analyzed for bisabolene and the aqueous layer was sampled for OD measurement and carbon utilization analysis (n=3, data shown as average ±s.d, representative from at least four independent experiments).

Much effort has been expended on metabolic engineering of common microbial host organisms such as *E. coli* and *S. cerevisiae* for simultaneous utilization of multiple carbon sources, such as glucose and xylose [20, 31]. The ability of *R. toruloides* to efficiently utilize multiple carbon sources, particularly hexose and pentose sugars combined with aromatic compounds, is something that even extensively engineered strains of *S. cerevisiae* and *E. coli* have been unable to accomplish. However, the decrease in glucose and xylose consumption rates in *R. toruloides* cultures grown on mixed sugars merits further investigation to determine if there is competitive sugar transport, catabolite repression or other mechanisms affecting the kinetics.

### Consumption of lignin-related monoaromatics

The ability of *R. toruloides* to consume *p*-coumaric acid indicates that it may also have the natural ability to consume other lignin-related aromatic compounds. To better understand this potential, *R. toruloides* was tested for its ability to consume compounds related to possible lignin degradation products derived from the H, G and S subunits. Compounds were tested with the same aromatic motif as the 4-hydroxyphenyl H units (*p*-coumaric acid and *p*-hydroxybenzoic acid), the 4-hydroxy-3-methoxyphenyl G units (ferulic acid and vanillic acid), as well as the non-substituted aromatic compound benzoic acid. Complete consumption of each substrate occurred within 72 hours in single carbon cultures, except for vanillic acid, which may have been toxic at the level tested (Fig. 6). We also tested a compound with the same aromatic motif as the 4- hydroxy-3,5-dimethoxypheny S units (sinapic acid), but the results were inconclusive due to apparent oxidation and precipitation of the substrate during the cultivation. Overall, these results indicate that *R. toruloides* has the metabolic potential to consume lignin-degradation products derived from depolymerization processes that produce compounds similar to those tested. This capability highlights the potential of *R. toruloides* to be used as a conversion host of monoaromatic lignin degradation products, a characteristic that will become more important as biomass deconstruction technologies advance to provide more extensive lignin depolymerization. However, lignin depolymerization technologies can produce a very diverse and heterogeneous mixture of aromatic and non-aromatic compounds, so although *R. toruloides* has the potential to convert certain compounds derived from depolymerized lignin, much work would need to be done to tailor both the structure of lignin and lignin-depolymerization strategies to bias the product range toward compounds that can be readily consumed by this organism. For example, one possible strategy to do this would be to engineer bioenergy feedstocks to generate lignin more heavily acylated with *p*-coumarate, which could then be readily liberated from the lignin under alkaline pretreatment conditions and converted into bioproducts [32].

**Fig. 6.**
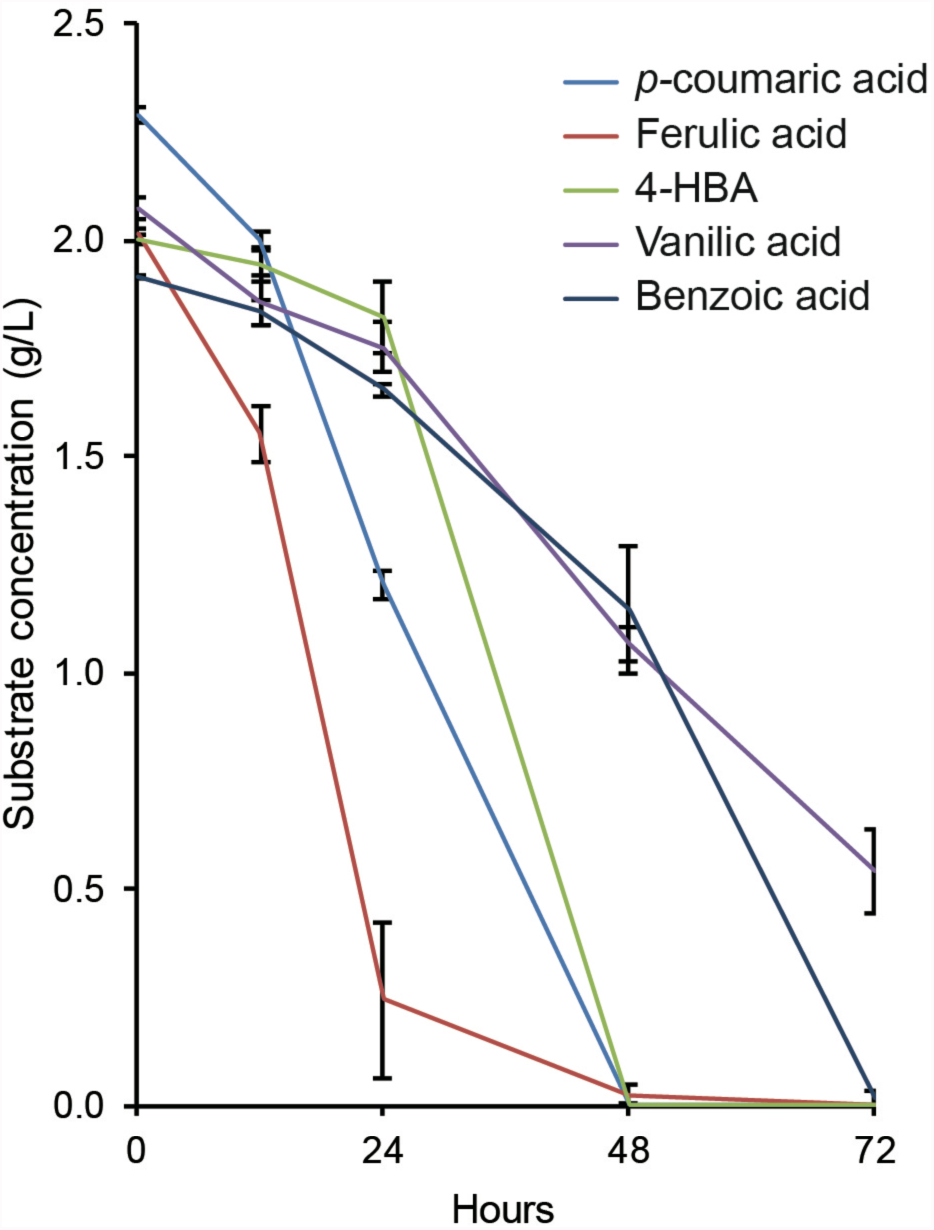
Utilization of several lignin-related aromatic compounds by *R. toruloides.* Carbon source utilization of *R. toruloides* grown in SD medium supplemented with 2 g/L of either *p*-coumaric acid, *p*-hydroxybenzoic acid (4-HBA), ferulic acid, vanillic acid, or benzoic acid (n=3, data shown as average ±s.d, from a single experiment).

### Bisabolene production and carbon source utilization in a corn stover hydrolysate pretreated with the novel biocompatible ionic liquid choline α-ketoglutarate

The performance of *R. toruloides* grown on purified substrates indicates that it may be an excellent biocatalyst for the conversion of deconstructed lignocellulose into valuable bioproducts. To test this premise, we examined how *R. toruloides* performs on substrates derived from actual lignocellulosic biomass. There are many technologies that have been developed to efficiently depolymerize biomass into intermediates suitable for microbial conversion, and those based on ionic liquid (IL) pretreatment and enzymatic saccharification have been demonstrated to be some of the most efficient and effective [33-35]. Recently, biocompatible ILs that do not inhibit commercial cellulase enzyme mixtures or microbial growth have been developed, enabling single-unit operation biomass pretreatment, saccharification, and fermentation, potentially reducing both CAPEX and OPEX in a biorefinery [36, 37]. Therefore, to test the performance of *R. toruloides* on a biomass hydrolysate, a corn stover hydrolysate containing glucose, xylose, and *p*-coumaric acid was generated using pretreatment with a novel biocompatible IL, choline α-ketoglutarate ([Ch][α-Kg]), followed by enzymatic saccharification (see methods for details). This IL falls into a recently developed class of ILs based on dicarboxylic acids [36]. Compositional and X-ray diffraction data, suggest that the IL pretreatment reduced recalcitrance to enzymatic saccharification by removing high amounts of lignin from the biomass and reducing the cellulose crystallinity (Table 3 and Additional file 1). *R. toruloides* was able to grow in the [Ch][α-Kg] hydrolysate, completely consume glucose, xylose, and *p*-coumaric acid, and produce 261 mg/L of bisabolene (Fig. 7A). In fact, it produced higher titers of bisabolene in the hydrolysate than it did in a control medium with matching concentrations of the IL, sugars, and *p*-coumaric acid (127 mg/L) (Fig. 7B).

**Table 3.** Chemical composition of dominant components in the dry corn stover before and after pretreatment.

|  | Solid recovery/ % | Glucan/ wt% | Xylan/ wt% | Lignin/ wt% |
| --- | --- | --- | --- | --- |
| Untreated Corn stover | / | 38.90±0.04 | 24.77±0.01 | 18.42±0.27 |
| Pretreated Corn stover | 60.0 | 48.35±0.06 | 28.92±0.05 | 13.12±0.03 |

Pretreatment conditions: 20% biomass loading, 80% [Ch][ α-Kg] (40 wt% in H<sub>2</sub>O), 120°C, 4 h.

**Fig. 7.**
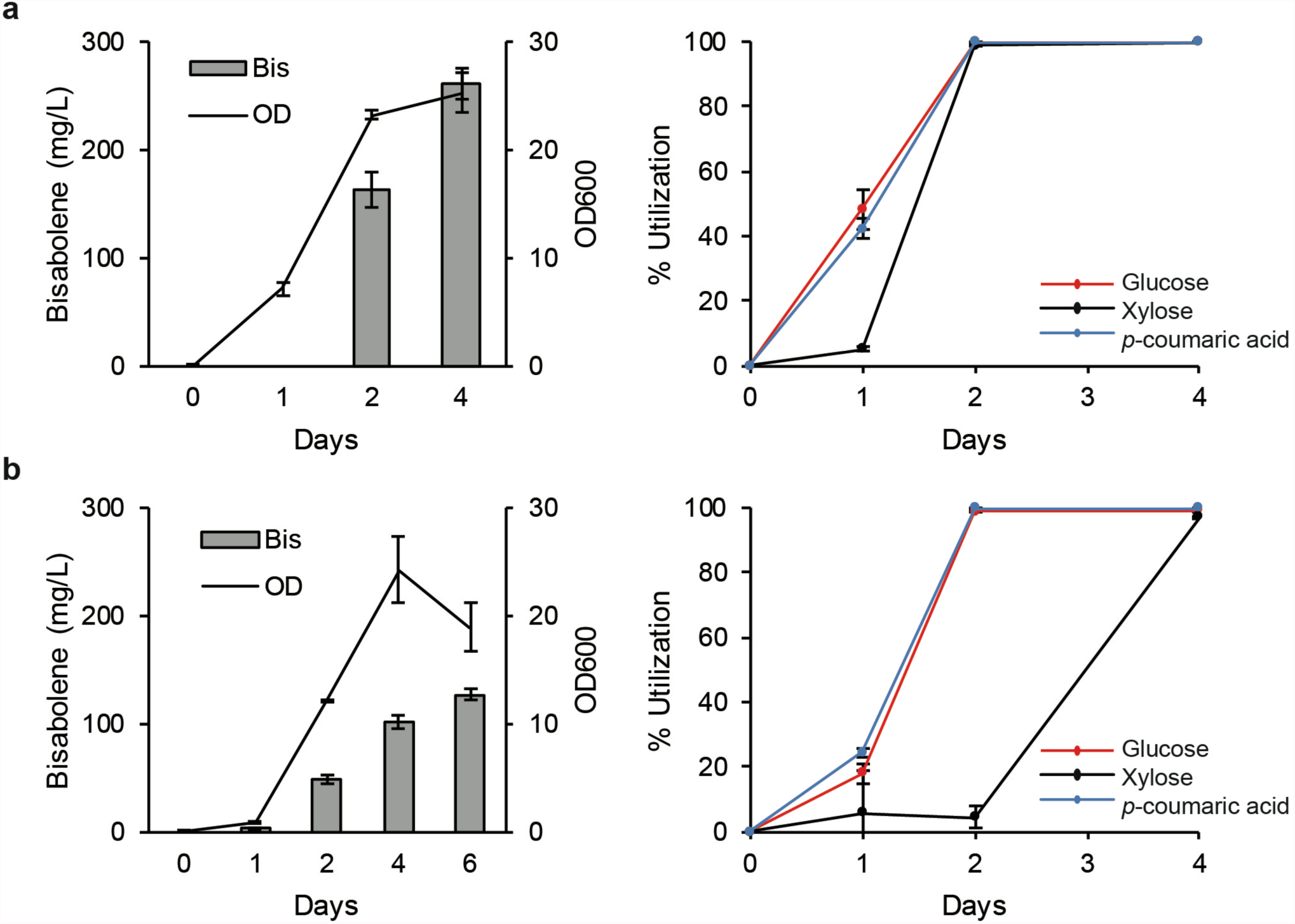
Conversion of biomass-derived glucose, xylose, and *p-*coumaric acid into bisabolene by *R. toruloides.* Bisabolene titers, growth, and carbon source utilization of strain BIS3 grown in (**a**) corn stover hydrolysate and (**b**) SD medium supplemented with individual components at the same concentration as those found in the corn stover hydrolysate: glucose (17.1 g/L), xylose (9.1 g/L), *p*- coumaric acid (383 mg/L), alphα-ketoglutarate (254 mM), and choline (586 mM). A low level of arabinose (0.98 g/L) was also detected in the hydrolysate and included in the control medium. Left panels: Lines represent ODs, bars represent bisabolene titers. Right panels: glucose (red), xylose (black), *p-*coumaric acid (blue). 5 mL cultures in test tubes were set up at a starting OD of 0.1 with a 20% dodecane overlay. At each time point, the dodecane layer was sampled and analyzed for bisabolene and the aqueous layer was sampled for OD measurement and carbon utilization analysis (n=3, data shown as average ±s.d, representative from at least two individual experiments).

### *R. toruloides* converts corn stover hydrolysate into bisabolene in high-gravity fed-batch bioreactors

In order to examine the spectrum of hydrolysates that *R. toruloides* can utilize and determine the impact of optimized cultivation conditions on bisabolene titers, a corn stover hydrolysate generated from an alkaline pretreatment was also tested. This pretreatment method generates very high concentrations of glucose and xylose, so it can be used for high-gravity fed-batch cultivation, which enables the addition of significantly more carbon than in the batch cultivations conducted with the [Ch][α-Kg] hydrolysate. The drawback to this approach is that lignin depolymerization products are removed during the process, so only glucose and xylose utilization can be examined. *R. toruloides* was cultivated in a controlled, high-gravity fed-batch bioreactor using the alkaline corn stover hydrolysate or a glucose-only control medium, and produced 680 mg/L and 521 mg/L bisabolene, respectively (Fig. 8A, B). The lower titer in the control may be due to the lower amount of sugars added to the cultivation (alkaline: 73.8 g/L and control: 61.5 g/L), resulting in a slightly lower dry cell weight (alkaline: 27 g/L and control: 25 g/L) and lower bisabolene production. It is interesting to note that *R. toruloides* produced higher titers of bisabolene in both hydrolysates versus their control media. In many instances, the opposite has been observed for organisms like *S. cerevisiae* and *E. coli*, further demonstrating the greater suitability of *R. toruloides* as a lignocellulosic conversion host.

**Fig. 8.**
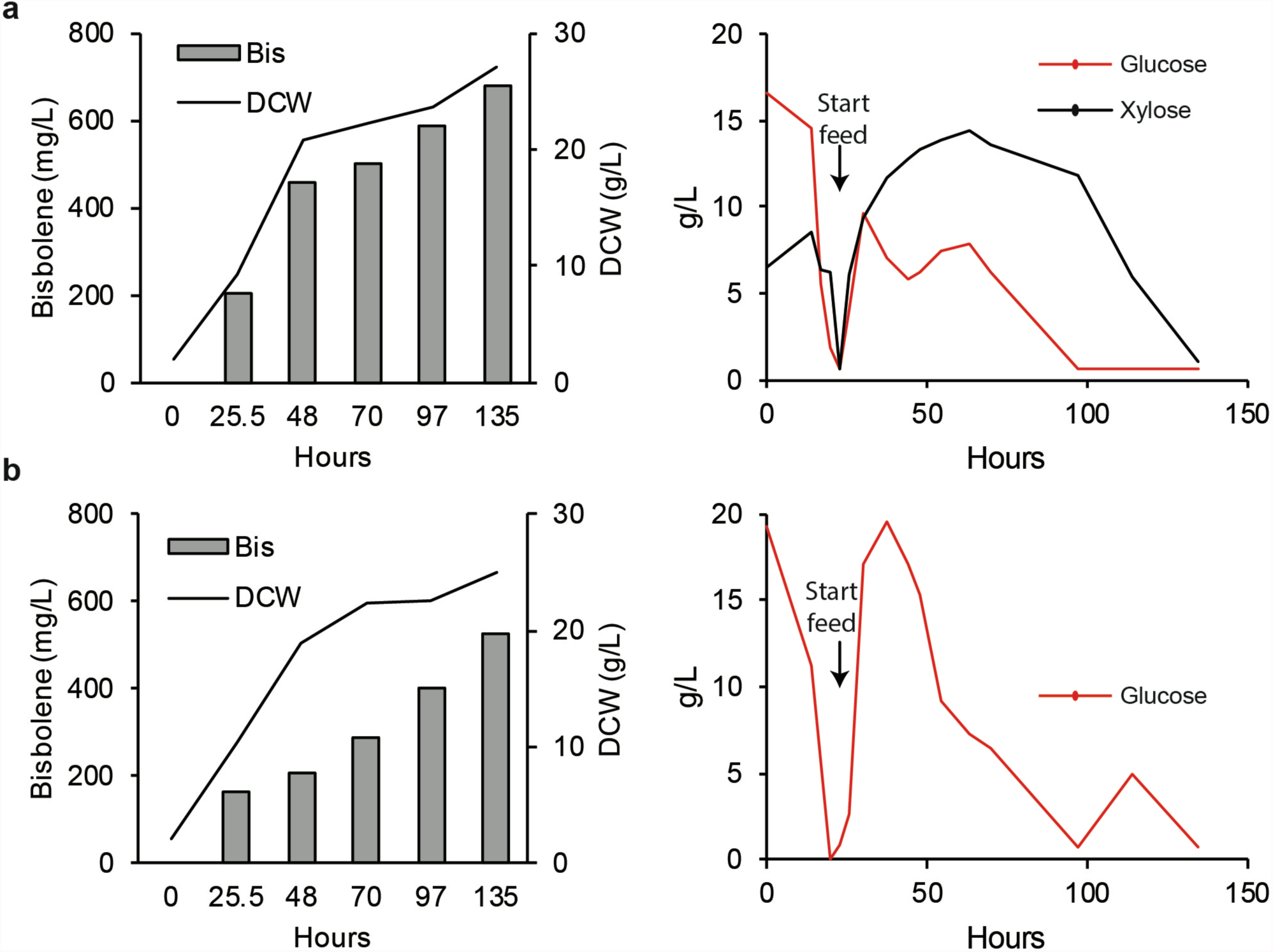
High-carbon fed-batch fermentation of *R. toruloides*. Bioreactor cultivation of strain BIS3 in (**a**) alkaline hydrolysate (**b**) SD medium with glucose. Left panels: Lines represent dry cell weight (DCW), bars represent bisabolene titers. Bisabolene titers were measured three times per time point, average value is shown. Right panels: Measured glucose (red) and xylose (black) concentration profiles. At each time point, 10 mL of the culture was sampled. After separation, the dodecane layer was used for bisabolene measurement. 5 mL of the aqueous layer was used for the measurement of DCW.

These results show that *R. toruloides* is amenable to high-carbon fed-batch fermentations, which is another important feature when considering organisms for use in industry. The bisabolene titer of 680 mg/L achieved in this study is impressive relative to titers reported for strains of *S. cerevisiae* and *E. coli* that have undergone extensive genetic engineering [24, 25, 28]. In addition, no significant reduction in the native pools of lipids or carotenoids was observed in the bisabolene-producing strains compared to wild type, suggesting that significant increases in titer can be achieved by further strain engineering to divert carbon flux away from these native molecules (both of which are derived from acetyl-CoA) toward bisabolene (Additional file 2).

## Conclusions

*Rhodosporidium toruloides* is emerging as a promising new production platform for the conversion of lignocellulose into biofuels and bioproducts. Much effort has focused on its oleaginous properties (high lipid proportions; >60% w/w of cell mass), and it has been engineered to produce several lipid derivatives [8, 38]. It has also been examined for its production of potentially valuable native carotenoids: β-carotene, torularhodin and torulene [39]. In this study, we demonstrate that this organism is a versatile production host that possesses many features critical to reducing CAPEX and OPEX in a biorefinery: 1) it can be used to make a variety of bioproducts, including non-native terpenes with biofuel and pharmaceutical applications, 2) heterologous production of bioproducts does not require inducers or antibiotics and is stable through multiple generations, 3) it can efficiently utilize components of both the polysaccharide and lignin fractions of inexpensive, carbon neutral, and renewable lignocellulosic feedstocks, 4) it is compatible with single-unit operation pretreatment, saccharification and fermentation bioprocessing configurations, and 5) bioproduct productivity is not inhibited in lignocellulosic hydrolysates. No other microbial production platform has been demonstrated to harbor all these properties, and *R. toruloides* sets a new standard for biotechnological applications that support a green economy.

## Abbreviations

ADS: amorphadiene synthase
ATMT: *Agrobacterium tumefaciens* mediated transformation
BIS: bisabolene synthase
CAPEX: capital expense
[Ch][α-Kg]: choline α-ketoglutarate
CrI: crystallinity index
CSM: complete supplemental mixture
DI: deionized
GC-MS: gas chromatography-mass spectrometry
HPLC: high performance liquid chromatography
IL: ionic liquid
LB: Luria Broth
LCA: life cyle analysis
OD: optical density
OPEX: operating expense
SD: synthetic defined
TEA: technoeconomic
XRD: X-ray diffraction
YNB: yeast nitrogen base
YPD: yeast peptone dextrose

## Declarations

### Ethics approval and consent to participate

Not applicable

### Consent for publication

Not applicable

### Availability of data and materials

All data generated or analysed during this study are included in this published article

### Competing interests

The authors declare no competing financial interests.

### Funding

This work was part of the DOE Joint BioEnergy Institute (http://www.jbei.org) supported by the U. S. Department of Energy, Office of Science, Office of Biological and Environmental Research, through contract DE-AC02-05CH11231 between Lawrence Berkeley National Laboratory and the U. S. Department of Energy.

### Author’s contributions

JMG and JMS conceived the project. JY and JK planned and performed experiments and analyzed data. JY, JK, and JMG wrote the manuscript. MI performed strain transformations. JS and TD performed the IL-pretreatment and hydrolysate preparation and analysis. MM, ERS and DT performed bioreactor experiments. AR performed experiments on consumption of lignin-related aromatics. EB performed LC-MS analysis. TP, KS, SS, JDK, BAS, SWS, JKM and APA edited the manuscript.

## Acknowledgments

This work was part of the DOE Joint BioEnergy Institute (http://www.jbei.org) supported by the U. S. Department of Energy, Office of Science, Office of Biological and Environmental Research, through contract DE-AC02-05CH11231 between Lawrence Berkeley National Laboratory and the U. S. Department of Energy. The United States Government retains and the publisher, by accepting the article for publication, acknowledges that the United States Government retains a non-exclusive, paid-up, irrevocable, world-wide license to publish or reproduce the published form of this manuscript, or allow others to do so, for United States Government purposes.

## Availability of data and materials

All data generated or analyzed during this study are included in this article and its additional files.

## Competing financial interests

The authors declare no competing financial interests.

## Additional files

### Additional file 1 (Additional file 1.pdf)

X-ray diffraction patterns and CrI (%) values of corn stover, both untreated and pretreated with [Ch][α-Kg] (40 wt% in H2O) at 120°C for 4 h pretreatment condition.

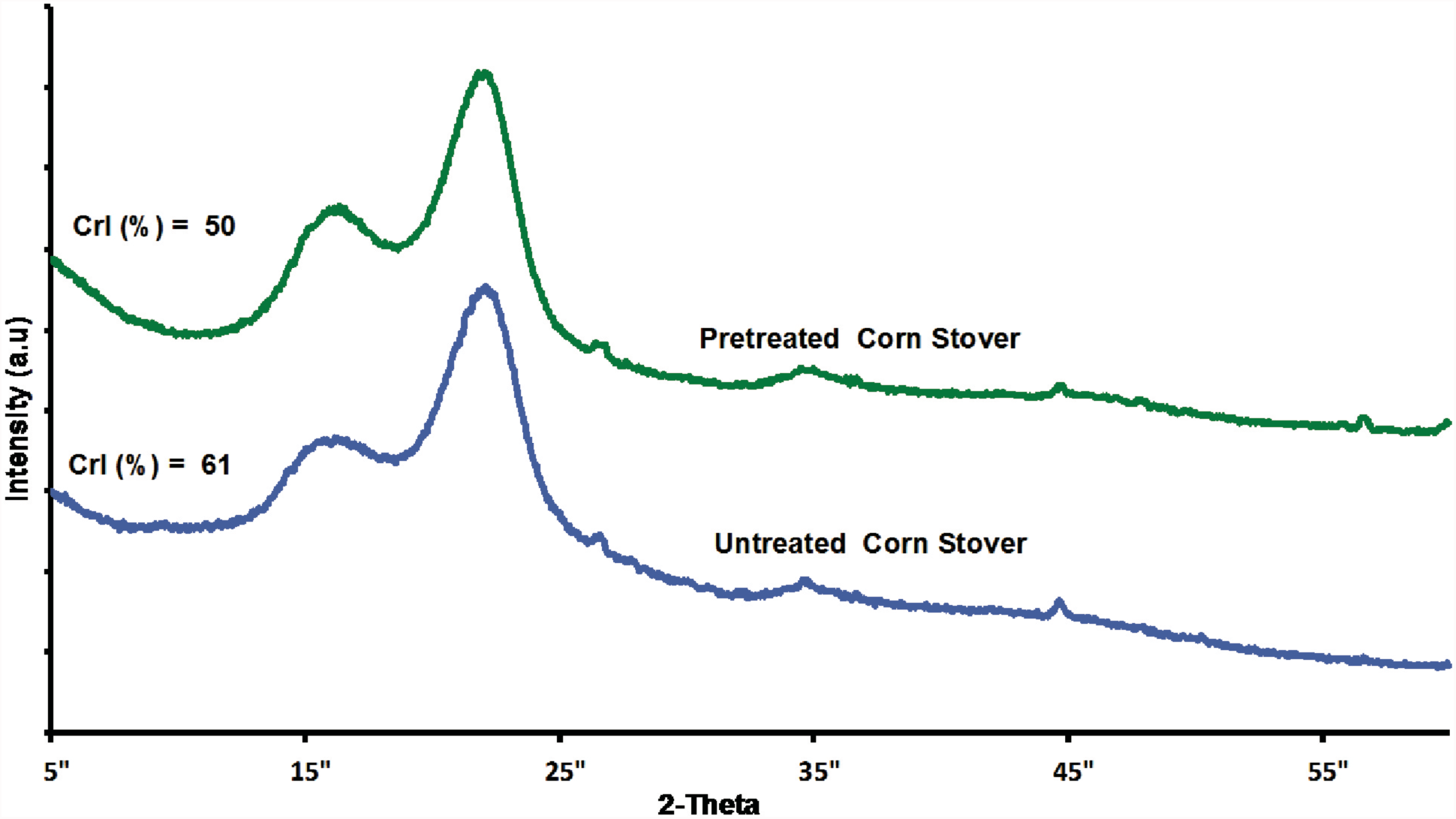

### Additional file 2 (Additional file 2.pdf)

Comparison of bisabolene titers, lipid content, and carotenoid levels between different BIS transformants. (n=3, data shown as average ±s.d, from a single experiment)

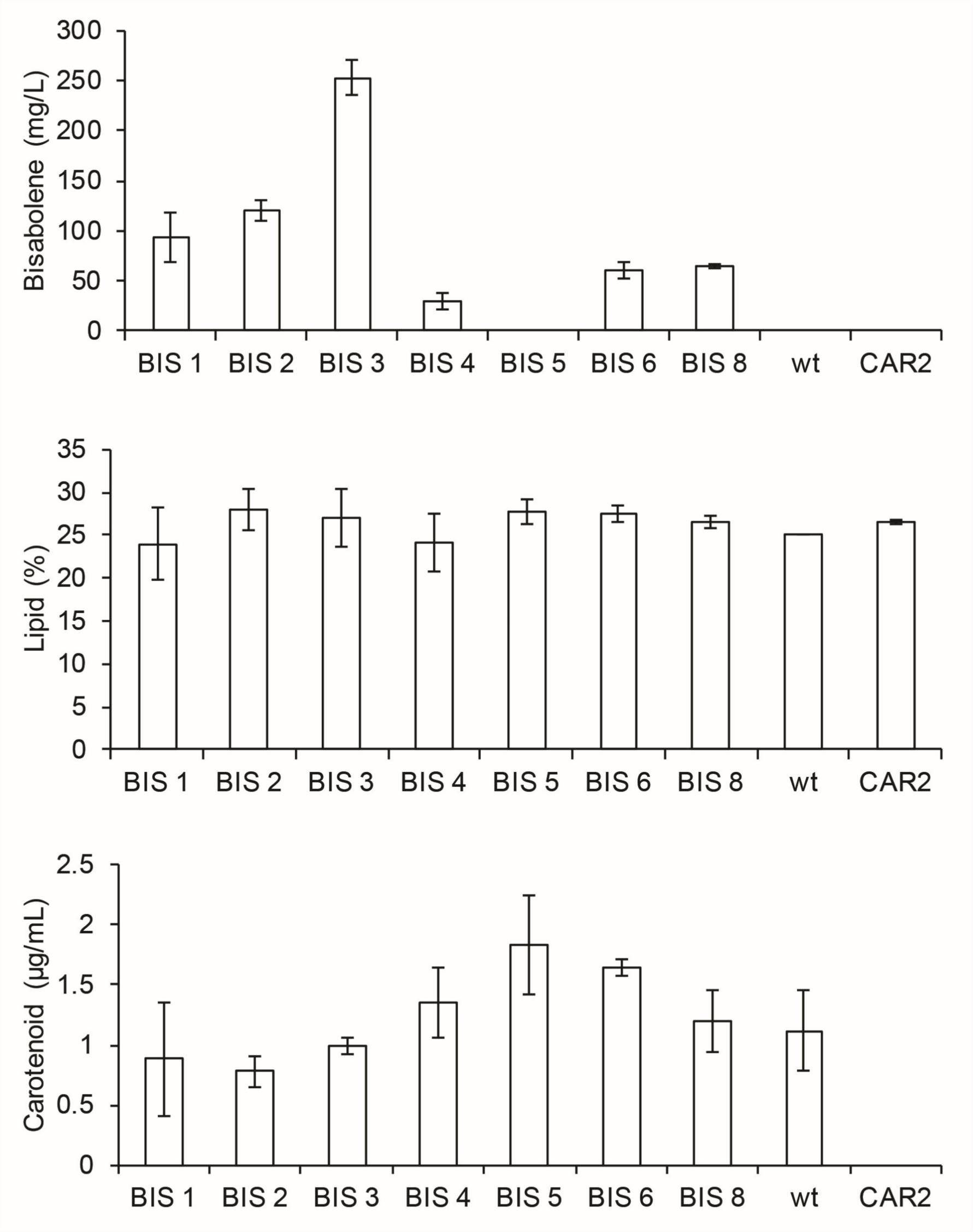

## References

1. Clark JH, Budarin V, Deswarte FEI, Hardy JJE, Kerton FM, Hunt AJ, Luque R, Macquarrie DJ, Milkowski K, Rodriguez A, Samuel O, Tavener SJ, White RJ, Wilson AJ. Green chemistry and the biorefinery: a partnership for a sustainable future. Green Chem. 2006, 8:853.

2. Himmel ME, Ding SY, Johnson DK, Adney WS, Nimlos MR, Brady JW, Foust TD. Biomass recalcitrance: Engineering plants and enzymes for biofuels production. Science 2007, 315:804–807.

3. Ragauskas AJ, Beckham GT, Biddy MJ, Chandra R, Chen F, Davis MF, Davison BH, Dixon RA, Gilna P, Keller M, Langan P, Naskar AK, Saddler JN, Tschaplinski TJ, Tuskan GA, Wymann CE. Lignin valorization: improving lignin processing in the biorefinery. Science 2014, 344:1246843.

4. Zaldivar J, Nielsen J, Olsson L. Fuel ethanol production from lignocellulose: a challenge for metabolic engineering and process integration. Appl Microbiol Biotechnol. 2001, 56:17–34.

5. Wiebe MG, Koivuranta K, Penttila M, Ruohonen L. Lipid production in batch and fed-batch cultures of Rhodosporidium toruloides from 5 and 6 carbon carbohydrates. BMC Biotechnol. 2012, 12.

6. Zhu ZW, Zhang SF, Liu HW, Shen HW, Lin XP, Yang F, Zhou YJJ, Jin GJ, Ye ML, Zou HF, Zhao ZBK. A multi-omic map of the lipid-producing yeast Rhodosporidium toruloides. Nat Commun. 2012, 3.

7. Ageitos JM, Vallejo JA, Veiga-Crespo P, Villa TG. Oily yeasts as oleaginous cell factories. Appl Microbiol Biotechnol. 2011, 90:1219–1227.

8. Zhang S, Skerker JM, Rutter CD, Maurer MJ, Arkin AP, Rao CV. Engineering Rhodosporidium toruloides for increased lipid production. Biotechnol Bioeng. 2016, 113:1056–1066.

9. Lin XP, Wang YN, Zhang SF, Zhu ZW, Zhou YJJ, Yang F, Sun WY, Wang XY, Zhao ZBK. Functional integration of multiple genes into the genome of the oleaginous yeast Rhodosporidium toruloides. FEMS Yeast Res. 2014, 14:547–555.

10. Liu YB, Koh CMJ, Sun LH, Hlaing MM, Du MG, Peng N, Ji LH. Characterization of glyceraldehyde-3-phosphate dehydrogenase gene RtGPD1 and development of genetic transformation method by dominant selection in oleaginous yeast Rhodosporidium toruloides. Appl Microbiol Biotechnol. 2013, 97:719–729.

11. Koh CMJ, Liu YB, Moehninsi, Du MG, Ji LH. Molecular characterization of KU70 and KU80 homologues and exploitation of a KU70-deficient mutant for improving gene deletion frequency in Rhodosporidium toruloides. BMC Microbiol. 2014, 14.

12. Hatfield RD, Marita JM, Frost K, Grabber J, Ralph J, Lu FC, Kim H. Grass lignin acylation: p-coumaroyl transferase activity and cell wall characteristics of C3 and C4 grasses. Planta 2009, 229:1253–1267.

13. Xu F, Sun J, Konda N, Shi J, Dutta T, Scown CD, Simmons BA, Singh S. Transforming biomass conversion with ionic liquids: process intensification and the development of a high-gravity, one-pot process for the production of cellulosic ethanol. Energy Environ Sci. 2016, 9:1042–1049.

14. Sun J, Konda N, Shi J, Parthasarathi R, Dutta T, Xu F, Scown CD, Simmons BA, Singh S. CO2 enabled process integration for the production of cellulosic ethanol using bionic liquids. Energy Environ Sci. 2016, 9:2822–2834.

15. Ham TS, Dmytriv Z, Plahar H, Chen J, Hillson NJ, Keasling JD. Design, implementation and practice of JBEI-ICE: an open source biological part registry platform and tools. Nucleic Acids Res. 2012, 40.

16. Abbott EP, Ianiri G, Castoria R, Idnurm A. Overcoming recalcitrant transformation and gene manipulation in Pucciniomycotina yeasts. Appl Microbiol Biotechnol. 2013, 97:283–295.

17. Ozaydin B, Burd H, Lee TS, Keasling JD. Carotenoid-based phenotypic screen of the yeast deletion collection reveals new genes with roles in isoprenoid production. Metab Eng. 2013, 15:174–183.

18. Sitepu IR, Garay LA, Sestric R, Levin D, Block DE, German JB, Boundy-Mills KL. Oleaginous yeasts for biodiesel: current and future trends in biology and production. Biotechnol Adv. 2014, 32:1336–1360.

19. Lee JJ, Chen L, Shi J, Trzcinski A, Chen WN. Metabolomic profiling of Rhodosporidium toruloides grown on glycerol for carotenoid production during different growth phases. J Agric Food Chem. 2014, 62:10203–10209.

20. Apel AR, Ouellet M, Szmidt-Middleton H, Keasling JD, Mukhopadhyay A. Evolved hexose transporter enhances xylose uptake and glucose/xylose co-utilization in Saccharomyces cerevisiae. Sci Rep. 2016, 6.

21. Park S BJ, Himmel ME, Parilla PA, Johnson DK. Research cellulose crystallinity index: measurement techniques and their impact on interpreting cellulase performance. Biotechnol Biofuels. 2010, 3.

22. Mansikkamäki P LM, Rissanen K. Structural changes of cellulose crystallites induced by mercerisation in different solvent systems; determined by powder X-ray diffraction method. Cellulose 2005, 12:233–242.

23. Maimone TJ, Baran PS. Modern synthetic efforts toward biologically active terpenes. Nat Chem Biol. 2007, 3:396–407.

24. Ro DK, Paradise EM, Ouellet M, Fisher KJ, Newman KL, Ndungu JM, Ho KA, Eachus RA, Ham TS, Kirby J, Chang MCY, Withers ST, Shiba Y, Saprong R, Keasling JD. Production of the antimalarial drug precursor artemisinic acid in engineered yeast. Nature 2006, 440:940–943.

25. Peralta-Yahya PP, Ouellet M, Chan R, Mukhopadhyay A, Keasling JD, Lee TS. Identification and microbial production of a terpene-based advanced biofuel. Nat Commun. 2011, 2.

26. Tyo KEJ, Ajikumar PK, Stephanopoulos G. Stabilized gene duplication enables long-term selection-free heterologous pathway expression. Nat Biotechnol. 2009, 27:760–U115.

27. Yin J, Wang H, Fu XZ, Gao X, Wu Q, Chen GQ. Effects of chromosomal gene copy number and locations on polyhydroxyalkanoate synthesis by Escherichia coli and Halomonas sp. Appl Microbiol Biotechnol. 2015, 99:5523–5534.

28. Paradise EM, Kirby J, Chan R, Keasling JD. Redirection of flux through the FPP branch-point in Saccharomyces cerevisiae by down-regulating squalene synthase. Biotechnol Bioeng. 2008, 100:371–378.

29. Kirby J, Nishimoto M, Chow RWN, Pasumarthi VN, Chan R, Chan LJG, Petzold CJ, Keasling JD. Use of Nonionic Surfactants for Improvement of Terpene Production in Saccharomyces cerevisiae. Appl Environ Microbiol. 2014, 80:6685–6693.

30. Jiang KK, Li LL, Long LK, Ding SJ. Comparison of alkali treatments for efficient release of p-coumaric acid and enzymatic saccharification of sorghum pith. Bioresour Technol. 2016, 207:1–10.

31. Groff D, Benke PI, Batth TS, Bokinsky G, Petzold CJ, Adams PD, Keasling JD. Supplementation of Intracellular XylR Leads to Coutilization of Hemicellulose Sugars. Appl Environ Microbiol. 2012, 78:2221–2229.

32. Rinaldi R, Jastrzebski R, Clough MT, Ralph J, Kennema M, Bruijnincx PCA, Weckhuysen BM. Paving the Way for Lignin Valorisation: Recent Advances in Bioengineering, Biorefining and Catalysis. Angew Chem. 2016, 55:8164–8215.

33. Li CL, Sun L, Simmons BA, Singh S. Comparing the Recalcitrance of Eucalyptus, Pine, and Switchgrass Using Ionic Liquid and Dilute Acid Pretreatments. BioEnergy Res. 2013, 6:14–23.

34. Shi J, George KW, Sun N, He W, Li CL, Stavila V, Keasling JD, Simmons BA, Lee TS, Singh S. Impact of Pretreatment Technologies on Saccharification and Isopentenol Fermentation of Mixed Lignocellulosic Feedstocks. BioEnergy Res. 2015, 8:1004–1013.

35. Singh S, Cheng G, Sathitsuksanoh N, Wu D, Varanasi P, George A, Balan V, Gao X, Kumar R, Dale BE, Wyman CE, Simmons BA. Comparison of different biomass pretreatment techniques and their impact on chemistry and structure. Front Energy Res. 2015, 2.

36. Liszka MJ, Kang A, Konda N, Tran K, Gladden JM, Singh S, Keasling JD, Scown CD, Lee TS, Simmons BA, Sale KL. Switchable ionic liquids based on di-carboxylic acids for one-pot conversion of biomass to an advanced biofuel. Green Chem. 2016, 18:4012–4021.

37. Sun J, Konda N, Shi J, Parthasarathi R, Dutta T, Xu F, Scown CD, Simmons BA, Singh S. CO2 enabled process integration for the production of cellulosic ethanol using bionic liquids. Energy Environ Sci. 2016:2822–2834.

38. Fillet S, Gibert J, Suarez B, Lara A, Ronchel C, Adrio JL. Fatty alcohols production by oleaginous yeast. J Ind Microbiol Biotechnol. 2015, 42:1463–1472.

39. Buzzini P, Innocenti M, Turchetti B, Libkind D, van Broock M, Mulinacci N. Carotenoid profiles of yeasts belonging to the genera Rhodotorula, Rhodosporidium, Sporobolomyces, and Sporidiobolus. Can J Microbiol. 2007, 53:1024–1031.

